# Detection of viral pathogens with multiplex Nanopore MinION sequencing: be careful with cross-talk

**DOI:** 10.1101/308262

**Authors:** Yifei Xu, Kuiama Lewandowski, Sheila Lumley, Steven Pullan, Richard Vipond, Miles Carroll, Dona Foster, Philippa C Matthews, Timothy Peto, Derrick Crook

**Author notes:** Correspondence address. Yifei Xu, Nuffield Department of Medicine, University of Oxford, John Radcliffe Hospital, Oxford OX3 9DU, United Kingdom.

## Abstract

Metagenomic sequencing with the Oxford Nanopore MinION sequencer offers potential for point-of-care testing of infectious diseases in clinical settings. To improve cost-effectiveness, multiplexing of several, barcoded samples upon a single flow cell will be required during sequencing. We generated a unique sequencing dataset to assess the extent and source of cross barcode contamination caused by multiplex MinION sequencing. Sequencing libraries for three different viruses, including influenza A, dengue and chikungunya, were prepared separately and sequenced on individual flow cells. We also pooled the respective libraries and performed multiplex sequencing. We identified 0.056% of total reads in the multiplex sequencing data that were assigned to incorrect barcodes. Chimeric reads were the predominant source of this error. Our findings highlight the need for careful filtering of multiplex sequencing data before downstream analysis, and the trade-off between sensitivity and specificity that applies to the barcode demultiplexing methods.

## Background

Metagenomic sequencing has the potential to allow unbiased identification of pathogens from a clinical sample. It holds the promise to serve as a single and universal assay for diagnostics of infectious diseases directly from samples without the need for *a priori* knowledge [1–3]. In addition to identification of pathogen species, broad and deep metagenomic sequence data could provide information relevant to determining treatment and prognosis, detecting outbreaks and tracking infection epidemiology [4–7]. Next-generation sequencing (NGS) platforms can produce a massive throughput of data at a modest cost, however, its application in clinical diagnostic and public health has been limited by complexity, slowness, and capital investment.

The MinION is a palm-size, real-time, single-molecule genome sequencer developed by Oxford Nanopore Technologies (ONT). The MinION’s compact size and real-time nature could facilitate the application of metagenomic sequencing in point-of-care testing for infectious diseases, as demonstrated by several proof-of-concept studies, including identification of Chikungunya (CHIKV), Ebola (EBOV), and hepatitis C virus (HCV) from human clinical blood samples without target enrichment [8], and detection of bacterial pathogens from urine samples [9] and respiratory samples, without the need for prior culture [10].

The data throughput of MinION has greatly increased since its release in 2015, with each consumable flow cell now generating between 10 and 20 Gb of DNA sequence data. This allows users to make more efficient use of the flow cell (and reduce cost) by multiplexing several samples in a single sequencing run. ONT has developed PCR-free barcode sets that allow multiplexing of up to 12 samples.

Detection of influenza A virus in multiple respiratory samples could be one diagnostic use of a multiplexed MinION sequencing assay. However, when sequencing directly from samples with a potential wide range of viral titres, it is important to be aware of the potential for cross sample contamination, both during library preparation and the bioinformatic barcode demultiplexing stage following sequencing. Here, we present a unique MinION sequencing dataset and results of investigation into the extent and source of cross barcode contamination in multiplex sequencing.

### Data Description

We used a ferret nasal wash sample infected with influenza A virus as an exemplar and also spiked two aliquots of negative nasal wash samples from uninfected ferret (pre-existing unused stocks from an unrelated study) with dengue and chikungunya viruses separately. Neither of these viruses are relevant for clinical diagnostics in respiratory samples, but act here as clear, distinct markers for the assessment of cross sample contamination. The sequencing libraries for each sample were prepared in parallel, along with a negative nasal wash control, barcoded, and sequenced individually. We then pooled an aliquot of the sequencing libraries and performed multiplex MinION sequencing. Reads from the four individual runs and the multiplex run were then analyzed to investigate the extent and source of cross sample contamination.

### Sample preparation

RNA was extracted, using the QIAamp viral RNA kit (Qiagen) according to the manufacturer’s instructions, from ferret nasal wash containing influenza A (H1N1) virus (A/California/04/2009) and a pool of negative nasal wash samples. Aliquots of negative sample extract were spiked with either dengue (DENV) (strain TC861HA, Genbank:MF576311) or CHIKV (strain S27, Genbank:MF580946.1) viral RNA from The National collection of Pathogenic Viruses (www.phe-culturecollections.org.uk). Samples were DNase treated using TURBO DNase (ThermoFisher Scientific) and purified using the RNA Clean & Concentrator™-5 kit (Zymo Research). cDNA was prepared and amplified using a Sequence-Independent-Single-Primer-Amplification method [8] modified as described previously [11]. Amplified cDNA was quantified using the Qubit dsDNA HS Assay Kit (ThermoFisher Scientific), and 1μg was used as input for each MinION library preparation, with the exception of the negative control where the entire sample was used.

### MinION library preparation and sequencing

Ligation Sequencing Kit 1D (SQK-LSK108) and Native Barcoding Kit 1D (EXP-NBD103) were used according to the ONT standard protocols, with the exception that only one barcode was included in each of the four library preparations. Each library was run on an individual flow cell and a fifth pooled library was made by combining the four individually barcoded libraries. Libraries were sequenced on R9.4 flow cells. The study design is shown in Fig. 1.

**Figure 1.**
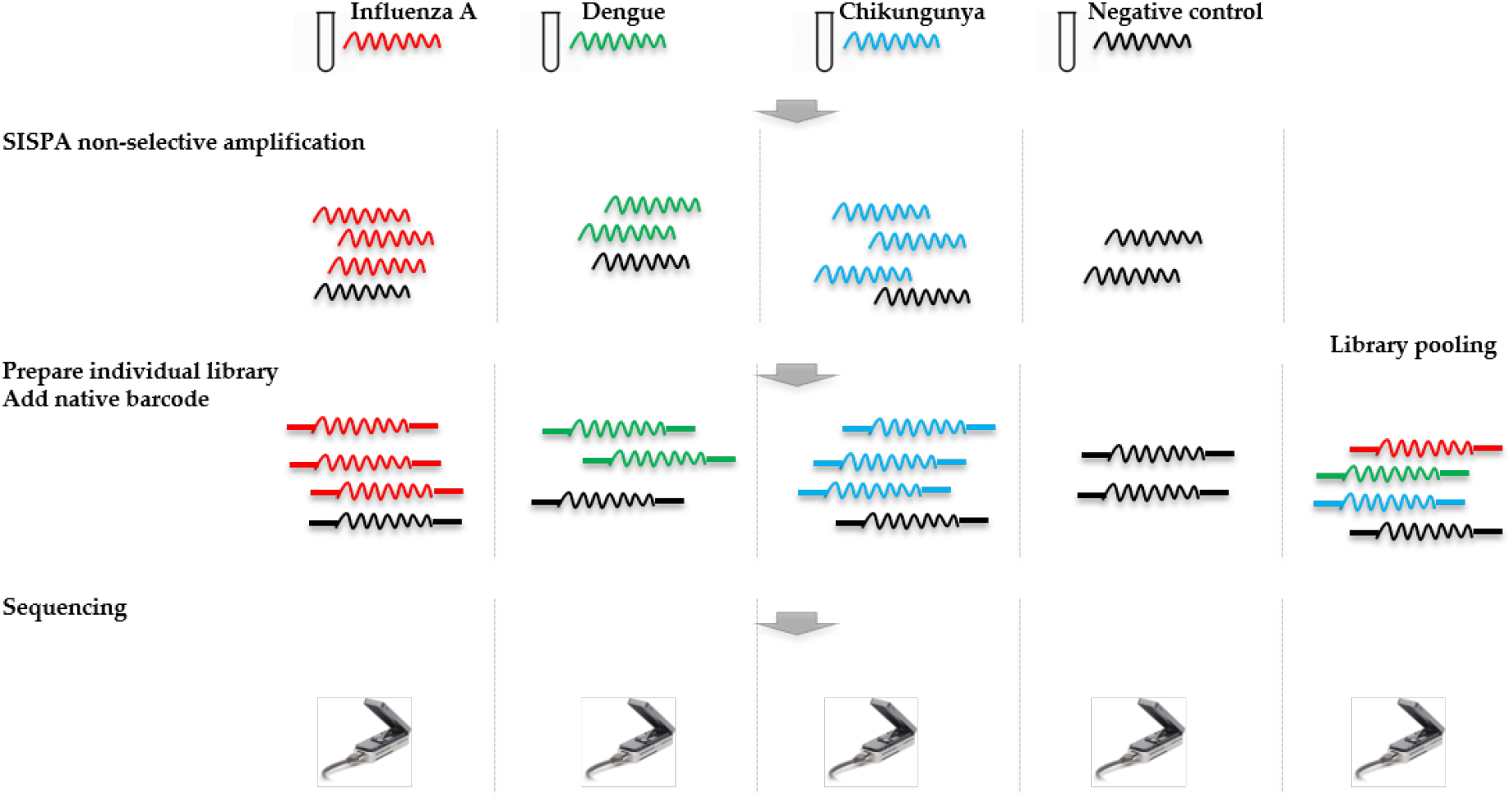
Overview of study design. RNA was extracted from four samples, including a ferret nasal wash sample infected with influenza A virus, two negative ferret nasal wash samples spiked with dengue and chikungunya viruses, and a negative ferret nasal wash control. cDNA was prepared and amplified using a Sequence-Independent-Single-Primer-Amplification method. The sequencing libraries for each sample were prepared in parallel, barcoded, and sequenced on individual flow cells. Multiplex sequencing was also performed by pooling the four individual libraries. Reads from the four individual runs and the multiplex run were analysed to assess the extent and source of cross barcode contamination in multiplex sequencing.

### Genomics analysis

Reads were basecalled using Albacore v2.1.7 (ONT) with barcode demultiplexing. Reads from each sequencing run were mapped to genomic sequences of each virus using Minimap2 [12]. The number of reads mapped to reference was counted using Pysam (https://github.com/pysam-developers/pysam). De novo assembly was performed using Canu v1.7 [13], and the resulting draft genome was polished using Nanopolish [14] with the signal-level data.

The barcode scores for each read were extracted from the “sequencing_summary.txt” file generated by Albacore. Two rounds of barcode demultiplexing were performed for the multiplex MinION sequencing data using Porechop (v0.2.2, https://github.com/rrwick/Porechop). Albacore’s output directory was used as input. First, we choose a threshold of 75 for “–middle_threshold” to examine internal adapter and filter chimera candidates. Next, we used the “–require_two_barcodes” option and set the threshold for barcode score as 70 to conduct stringent barcode binning. Read current signals were extracted from FAST5 file using ONT fast5 API (https://github.com/nanoporetech/ont_fast5_api) and plotted by using ggplot2 implemented in R (https://www.r-project.org/) for a comparison of chimeric and non-chimeric reads.

## Results

### MinION sequencing data and assembly of viral genomes

The throughput of each MinION sequencing run varied due to differences in running time. A maximum number of ~2.4M reads was achieved by the multiplexed sequencing run and the individual CHIKV run, due to longer running times (Table S1). Reads from the spiked virus accounted for 96% of the data in the individual CHIKV and DENV sequencing runs, and 78% for the flu sample (Table 1). The percentage of viral reads within each barcoded sample in the multiplex sequencing data is close to that in the individually run sample data (Table 2). Each viral genome had an ultra-high mean depth of coverage in the individual and multiplex sequencing data, and *de novo* assembly was able to recover nearly complete genomes for all three viruses with 99.9% identities compared to the GenBank reference.

**Table 1.**
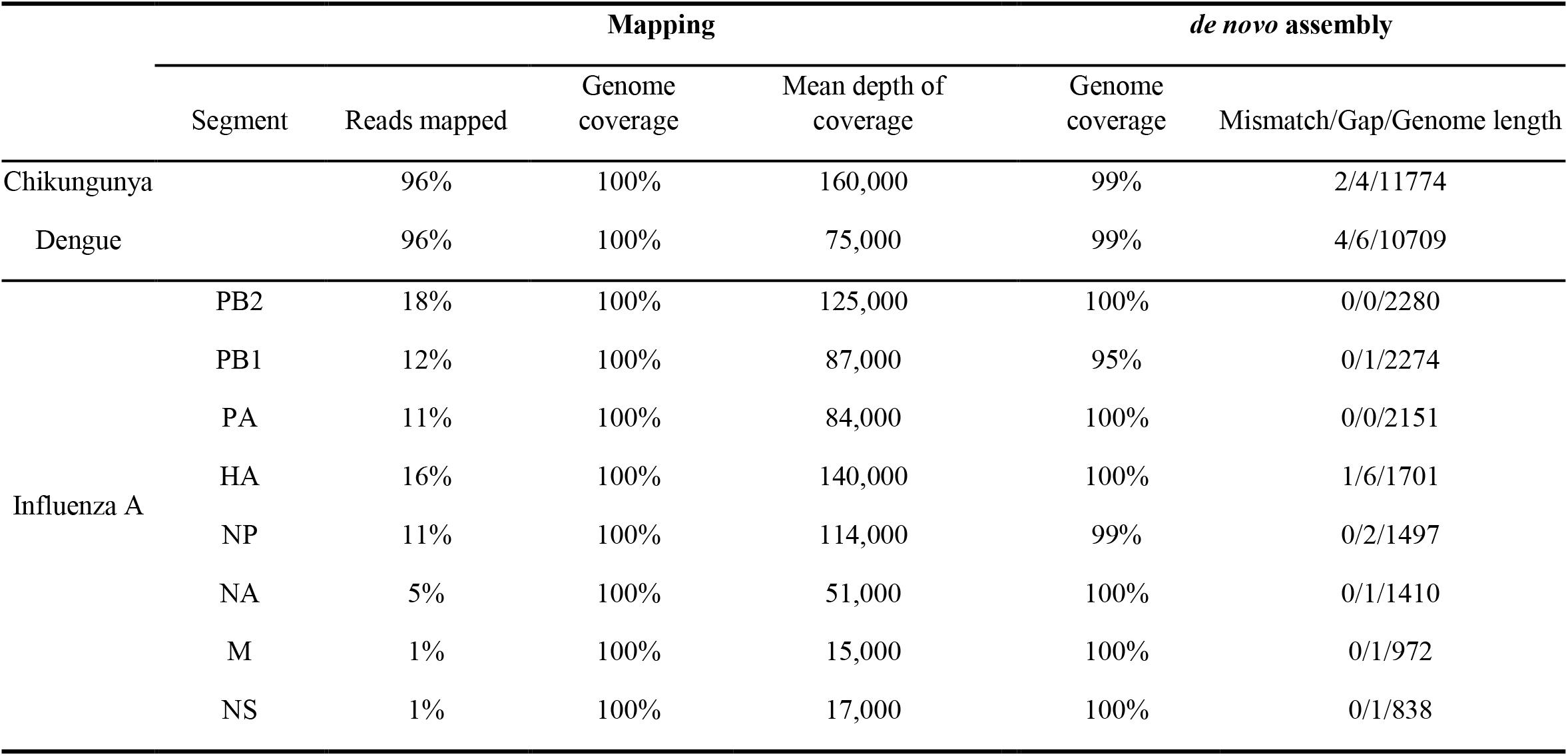
Summary of mapping and *de novo* assembly results for data from MinION sequencing of individual libraries. The eight gene segments of influenza A virus are PB2, polymerase basic 2; PB1, polymerase basic 1; PA, polymerase acidic; HA, hemagglutinin; NP, nucleoprotein; NA, neuraminidase; M, matrix; NS, non-structural.

**Table 2.**
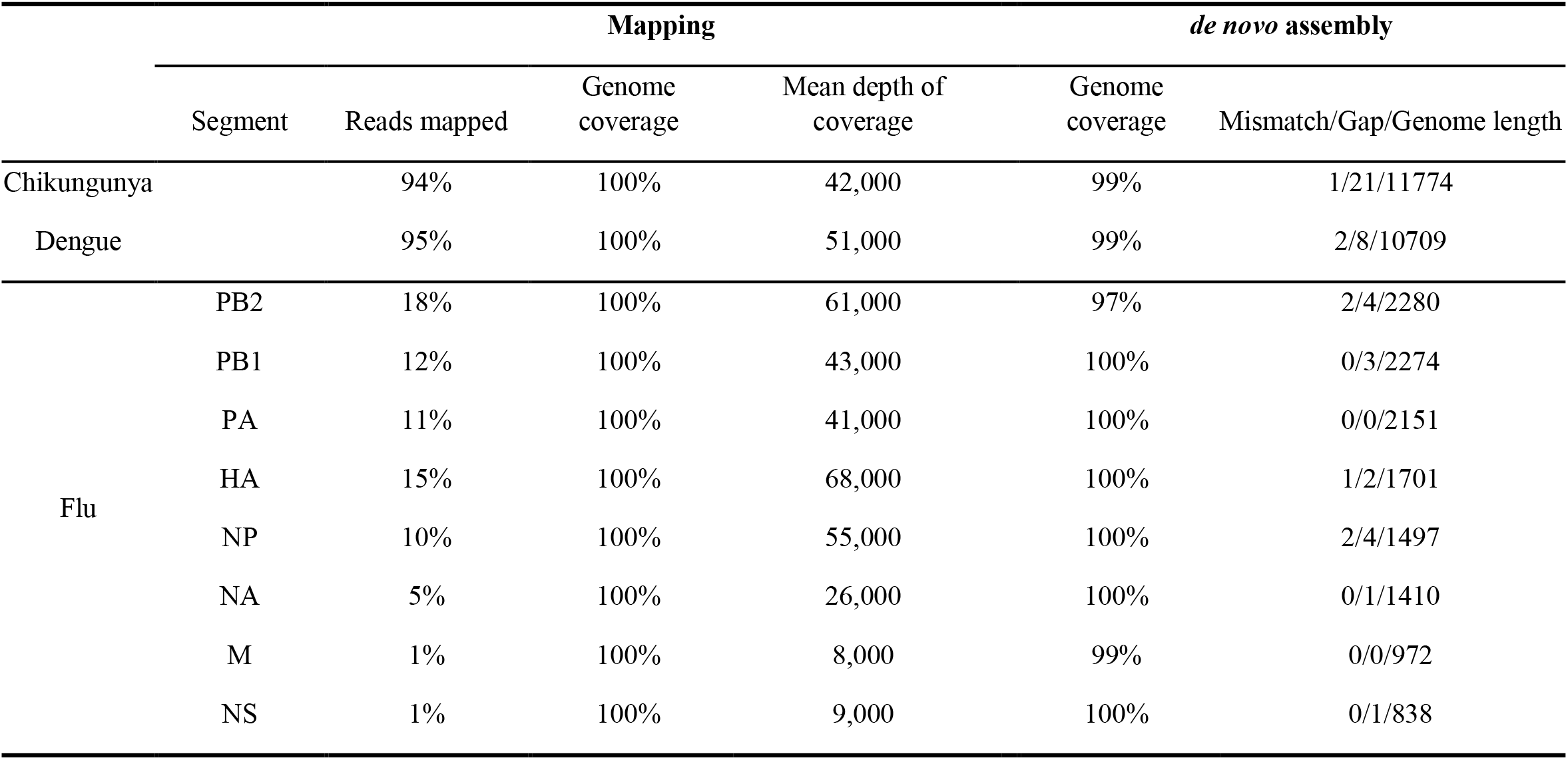
Summary of mapping and *de novo* assembly results for data from multiplex MinION sequencing. The eight gene segments of influenza A virus are PB2, polymerase basic 2; PB1, polymerase basic 1; PA, polymerase acidic; HA, hemagglutinin; NP, nucleoprotein; NA, neuraminidase; M, matrix; NS, non-structural.

|  |  | Mapping |  |  | *de novo* assembly |  |
| --- | --- | --- | --- | --- | --- | --- |
|  | Segment | Reads mapped | Genome coverage | Mean depth of coverage | Genome coverage | Mismatch/Gap/Genome length |
| Chikungunya |  | 94% | 100% | 42,000 | 99% | 1/21/11774 |
| Dengue |  | 95% | 100% | 51,000 | 99% | 2/8/10709 |
| Flu | PB2 | 18% | 100% | 61,000 | 97% | 2/4/2280 |
|  | PB1 | 12% | 100% | 43,000 | 100% | 0/3/2274 |
|  | PA | 11% | 100% | 41,000 | 100% | 0/0/2151 |
|  | HA | 15% | 100% | 68,000 | 100% | 1/2/1701 |
|  | NP | 10% | 100% | 55,000 | 100% | 2/4/1497 |
|  | NA | 5% | 100% | 26,000 | 100% | 0/1/1410 |
|  | M | 1% | 100% | 8,000 | 99% | 0/0/972 |
|  | NS | 1% | 100% | 9,000 | 100% | 0/1/838 |

### Extent and source of cross-sample contamination

Each sample was barcoded, and sequenced both individually and multiplexed, which allowed us to examine the performance of barcode demultiplexing of Albacore. In the individually sequenced sample data we would expect only a single native barcode to be present. For CHIKV (barcode NB01), DENV (NB09) and FLU-A (NB10) individual sequencing runs, we found that 86, 109, and 17 reads respectively were assigned to barcode bins not expected to be present in the library (representing 0.0036%, 0.0129%, and 0.001% of total reads). In the multiplex sequencing data, 41 reads (0.0016%) were assigned to barcodes not included in the experiments (i.e. a barcode other than NB01, NB05, NB09 or NB10). We defined these as mis-assigned reads (Figure 2a).

**Figure 2.**
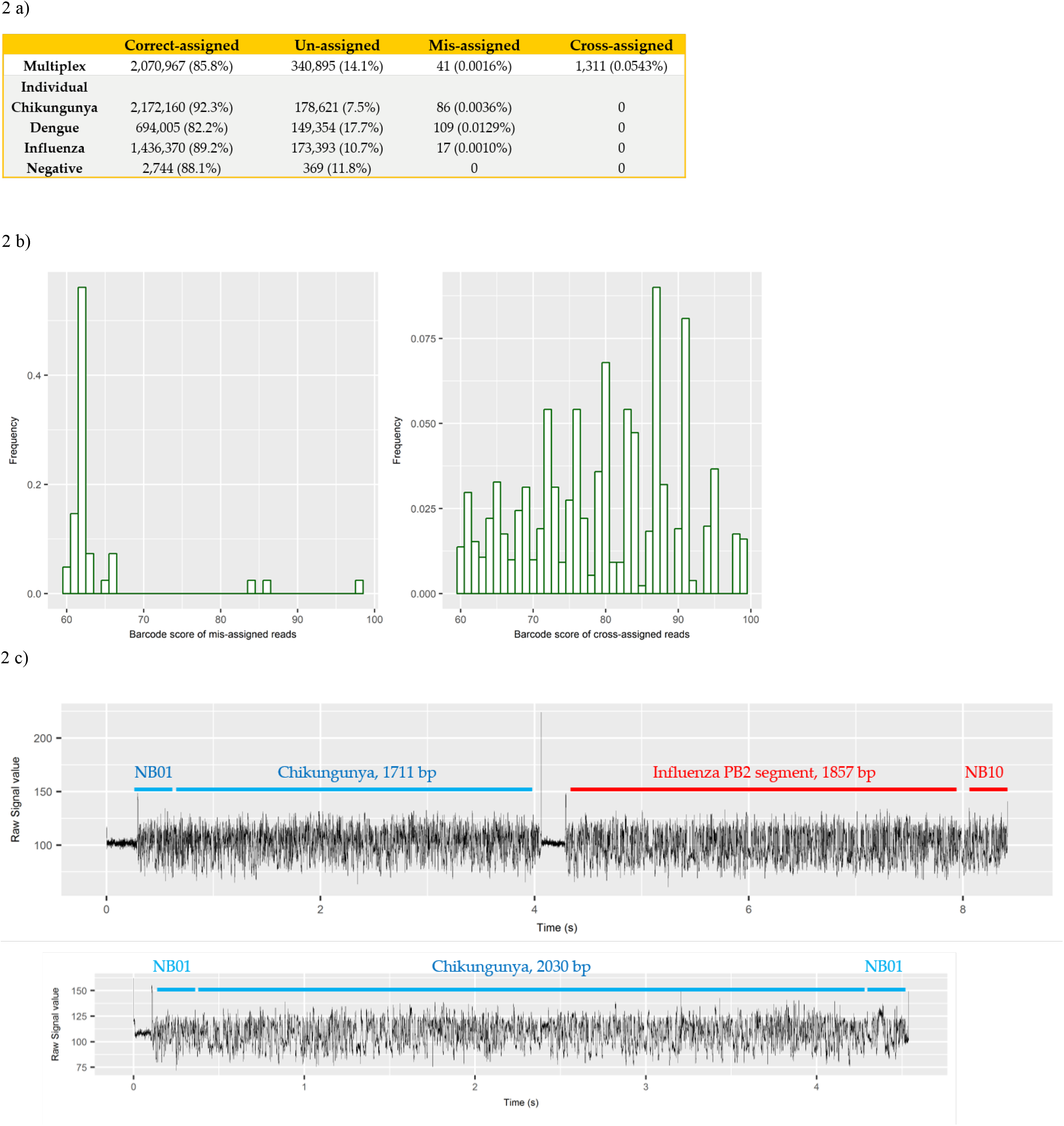
a) summary of number and percentage of reads correctly assigned, unassigned, mis-assigned, and cross-assigned in each sequencing run. Un-assigned refers to reads that cannot be assigned to any bins by Albacore due to a barcode score less than 60, mis-assigned refers to reads that were assigned to barcode bins not included in this experiment, and cross-assigned refers to reads that were assigned to the incorrect barcode bins; b) distribution of barcode scores reported by Albacore for mis-assigned reads and cross-assigned reads in the multiplex sequencing data; c) comparison of raw signal of a chimeric and a correctly assigned read. The signal of chimeric read possesses a stall signal and a huge spike signal in the middle of the read.

To examine potential laboratory contamination in sequencing library preparation, we mapped all reads from each individual run against the genomic sequences of all three viruses. No read was found to originate from a genome prepared in a different library, suggesting no *in vitro* contamination. The multiplex sequencing library was prepared by pooling the individual, non-contaminated libraries after the ligation of both barcode and adapter. However, mapping results show 1311 (0.0543%) reads mapped to the incorrect target genome, implying that they were cross-assigned to the wrong barcode bins (later referred to as ‘cross-assigned reads’), despite the fact that the multiplexed sequencing library was pooled with individual libraries showed no cross-assigned reads at all. We hypothesized that mis-assigned and cross-assigned reads were due to a low barcode score, and investigated the barcode scores of these reads. Most of the mis-assigned reads had a barcode score <70, however, cross-assigned reads had more diverse scores ranging from 60 to nearly 100 (Fig. 2b). This result suggested that mis-assigned and cross-assigned reads originate from different sources. We blasted the cross-assigned reads to a small database comprising the genomic sequences of the three viruses included in this study, and demonstrated that 1075/1311 (82%) of cross-assigned reads could be aligned to more than one viral genome or to distinct regions within the same genome, suggesting they are chimeras.

To confirm this observation, we investigated the raw current signals of a few cross-assigned reads compared to those of correctly assigned reads (Fig. 2c). The current signals of a correctly assigned read usually include: (i) an open pore signal of high current representing the time that the sequencing pore changes from one adapter to another, (ii) a stall signal, referring to the period of time that a DNA sequence is in the pore but yet to move, and (iii) the signal trace of DNA sequencing. In contrast, a chimeric read possesses a stall signal and a huge spike signal in the middle of the read. Chimeric reads can possess two different barcode sequences at the start and end, thus confusing assignment of a barcode bin. In addition, we also observed that a small number of reads possessed a 24 nucleotide sequence that is identical to a barcode sequence that is not expected to be present in the library. Taken together, these data demonstrate three categories of error that contribute to cross sample contamination in our dataset: (i) chimeric reads (account for ~80% of all cross-assigned reads); (ii) reads with low barcode score (account for <20% of all cross-assigned reads); (iii) reads possessing sequence that is 100% identical to an incorrect barcode sequence (infrequent).

In order to improve the quality of our final dataset, we explored the impact of different barcode demultiplexing approaches to remove cross-assigned reads (Fig. 3). Filtering of the reads that possess an internal adapter can remove 90% of the cross-assigned reads and lost 24% of total reads. We also tried a more stringent filtering scheme that required two barcodes (one each at the start and end of the read) to make an assignment. This approach removed all but two cross-assigned reads, but lost 56% of total reads.

**Figure 3.**
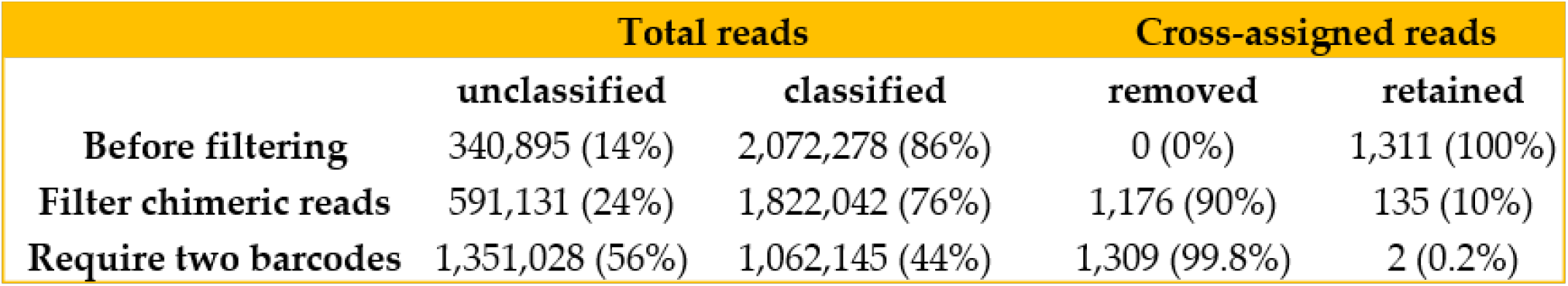
Removal of cross-assigned reads and loss of total sequencing data by two filtering approaches using Porechop. The first approach examines internal adapter and filter chimera candidates that possess adapter in the middle of the read. The second approach requires two barcodes at the start and end of each read. Increasing stringency for barcode demultiplexing occurs at the expense of reducing total number of reads.

## Discussion

The ultimate objective of our research is to develop a nanopore metagenomic sequencing based diagnostic assay that enables point-of-care testing for infectious diseases. Multiplex sequencing offers the opportunity to improve scalability and cut cost, however, cross sample contamination can lead to errors in the data and false interpretation of the results. Cross sample contamination has been observed before [15], but it has not previously been possible to separate library preparation from sequencing as the cause of chimeric reads. In this experiment, we pooled clean libraries and performed multiplex MinION sequencing in order to investigate the extent and source of cross-barcode contamination. We identified 0.056% of total reads were cross-assigned to the incorrect barcode bins, which is comparable to those reported for Illumina sequencing platforms from different studies (between 0.06% and 0. 25%) [16–18]. Our results showed that chimeric reads are the predominant source of cross-barcode assignment errors. Chimeric reads in this dataset could only have been formed during sequencing rather than library preparation, as they were completely absent in the sequencing data of individual libraries, and the only further processing step was to mix the final sequencing libraries prior to loading. We hypothesize that the current algorithm implemented in Albacore cannot recognize the short dissociation between DNA sequences that run concurrently through the nanopore, thereby concatenating more than one sequence into the same Fast5 file.

In summary, our study demonstrated that chimeric reads are the predominant source of cross barcode assignment errors in multiplex MinION sequencing. It highlights the need for careful filtering of multiplex MinION sequencing data before downstream analysis, and the trade-off between sensitivity and specificity that applies to the barcode demultiplexing methods.

## Author contributions

All authors designed the study. S.P., K. L., S. L., and Y.X. conducted MinION sequencing. Y.X. analyzed data. All authors participated in interpreting the results and writing the manuscript. All authors read and approved the final version of this manuscript.

## Acknowledgments

The authors would like to thank Dr. Anthony Marriott (Public Health England) for providing ferret nasal aspirates.

---

**Table S1.** Summary of statistics of MinION sequencing data.

|  | Virus | Barcode | SISPA concentration (ng/µl) | Number of reads | Total length (bp) |
| --- | --- | --- | --- | --- | --- |
| Individual sequencing | chikungunya | NB01 | 88.6 | 2,350,867 | 2,319,958,089 |
|  | dengue | NB09 | 70.2 | 843,468 | 965,165,841 |
|  | influenza A virus | NB10 | 49.4 | 1,609,780 | 1,749,616,221 |
|  | negative control | NB05 | 0.712 | 3,113 | 3,475,147 |
| Mutliplex sequencing | chikungunya | NB01 | - | 639,986 | 632,952,453 |
|  | dengue | NB09 | - | 580,183 | 666,239,544 |
|  | influenza A virus | NB10 | - | 823,427 | 881,448,974 |
|  | negative control | NB05 | - | 28,682 | 29,967,359 |
|  | unclassified |  |  | 340,895 | 360,996,367 |

